# Globally elevating the AGE clearance receptor, OST48, does not protect against the development of diabetic kidney disease, despite improving insulin secretion

**DOI:** 10.1101/710558

**Authors:** Aowen Zhuang, Felicia YT Yap, Domenica McCarthy, Sherman S. Leung, Sally A. Penfold, Chris Leung, Karly C. Sourris, Brooke E. Harcourt, Vicki Thallas-Bonke, Melinda T Coughlan, Benjamin L Schulz, Josephine M Forbes

**Affiliations:** Glycation and Diabetes Complications, Mater Research Institute – The University of Queensland, Translational Research Institute, Woolloongabba, Australia; School of Medicine, University of Queensland, St Lucia, Australia; Baker Heart and Diabetes Institute, Melbourne, Australia; Department of Immunology, Central and Eastern Clinical School, AMREP Precinct, Monash University, Australia; School of Biomedical Sciences, The University of Queensland, Brisbane, Australia; Department of Medicine, University of Melbourne, Austin Hospital, Heidelberg, Australia; Hormone and Obesity Research, Murdoch Childrens Research Institute, Melbourne, Australia; Department of Diabetes, Central Clinical School, Monash University, Melbourne, Australia; School of Chemistry and Molecular Biosciences, University of Queensland, St Lucia, Australia; Mater Clinical School, University of Queensland, St Lucia, Australia

**Author notes:** These authors contributed equally to this manuscript (Benjamin L Schulz, Josephine M Forbes). Contact Information Professor Josephine Forbes, Program Leader, Cardiometabolic Disease, Mater Research Institute – UQ, 37 Kent Street, Woolloongabba, QLD, 4102, Australia. Financial Support: This work has been supported through the funding bodies of the National Health Medical Research Council (NHMRC) Australia, Juvenile Diabetes Research Foundation (JDRF), Kidney Health Australia (KHA) and the Mater Research Foundation.

**Keywords:** advanced glycation end-product receptor 1 (AGE-R1), advanced glycation end-products (AGEs), diabetic nephropathy, diabetes

## Abstract

The accumulation of advanced glycation end products (AGEs) have been implicated in the development and progression of diabetic kidney disease (DKD). There has been interest in investigating the potential of AGE clearance receptors, such as oligosaccharyltransferase-48kDa subunit (OST48) to prevent the detrimental effects of excess AGE accumulation seen in the diabetic kidney. Here the objective of the study was to increase the expression of OST48 to examine if this slowed the development of DKD by facilitating the clearance of AGEs. Groups of 8-week-old heterozygous knock-in male mice (n=9-12/group) over-expressing the gene encoding for OST48, dolichyl-diphosphooligosaccharide-protein glycosyltransferase (*DDOST*+/-) and litter mate controls were randomised to either (i) no diabetes or (ii) diabetes induced via multiple low-dose streptozotocin and followed for 24 weeks. By the study end, global over expression of OST48 increased glomerular OST48. This facilitated greater renal excretion of AGEs but did not affect circulating or renal AGE concentrations. Diabetes resulted in kidney damage including lower glomerular filtration rate, albuminuria, glomerulosclerosis and tubulointerstitial fibrosis. In diabetic mice, tubulointerstitial fibrosis was further exacerbated by global increases in OST48. There was significantly insulin effectiveness, increased acute insulin secretion, fasting insulin concentrations and AUC_insulin_ observed during glucose tolerance testing in diabetic mice with global elevations in OST48 when compared to diabetic wild-type littermates. Overall, this study suggested that despite facilitating urinary-renal AGE clearance, there were no benefits observed on kidney functional and structural parameters in diabetes afforded by globally increasing OST48 expression. However, the improvements in insulin secretion seen in diabetic mice with global over-expression of OST48 and their dissociation from effects on kidney function warrant future investigation.

## Introduction

Currently, the world is faced with a pandemic of both type 1 and type 2 diabetes (T1D and T2D)^1,2^, defined by persistent hyperglycaemia, which is a predominant factor in the development of concomitant complications^3,4^. A major complication, diabetic kidney disease (DKD) is a worldwide health concern and is an important risk factor for end-stage renal disease (ESRD)^5^ and cardiovascular disease^3,6^. Understanding the pathophysiology of DKD development and progression is therefore, a key challenge to reduce the burden of diabetic complications. The early clinical presentation of DKD is characterised by hyperfiltration, progressive proteinuria and associative glomerular injury accompanied by tubulointerstitial fibrosis and in the later stages, a steady progressive decline in renal function^7^. Unfortunately, despite optimal clinical management, involving both glycaemic^8,9^ and blood pressure control^10^, including inhibitors of the renin-angiotensin-aldosterone system^11,12^, it is only possible to achieve a 30% improvement in declining kidney function in DKD. Despite intervention, many individuals reach end stage disease requiring renal replacement therapy or die prematurely from a cardiovascular event^13^. Hence new therapies to combat DKD are urgently required.

AGE accumulation, specifically the deposition of *N*-(carboxymethyl)lysine (CML) within tissues is a pathological mediator in DKD^14,15^. It is postulated that under physiological conditions, OST48 may facilitate AGE clearance into the urine^16^ and that this function is impaired during the development and progression of DKD^17^ resulting in pathological accumulation of AGE at sites such as the kidney. OST48 (also known as AGER1) dually functions as an extracellular AGE binding protein^18^ and as a 48kDa subunit that functions as part of the oligosaccharyltransferase complex, which mediates the transfer of high-mannose oligosaccharides to asparagine residues within the lumen of the rough ER. OST48 gene and protein expression are decreased by diets abundantly rich in AGEs^19^ and by diabetes^20,21^. In addition, there are associations between lower OST48 levels in circulating immune cells and progressive diabetic nephropathy in a small cohort of patients with type 1 diabetes^22,23^ and with impaired insulin sensitivity patients with in type 2 diabetes^24,25^. To date, there has been one, *in vivo* investigation using untargeted over-expression of OST48. This study in elderly mice (>500 days old) demonstrated improvements in longevity, insulin sensitivity and resistance to balloon injury in blood vessels^26^. However, there are no studies where the efficacy of increasing OST48 expression to facilitate AGE clearance has been tested in the development of kidney disease including DKD.

A SNP array from the FinnDiane population study (*n* = 2719) inferred that there was an association of the OST48 gene, *DDOST* (rs2170336), with nephropathy development in individuals with T1D^27^. Additionally, a study in T1D patients identified that elevated serum AGEs were associated with increased OST48 mRNA in circulating mononuclear cells^22^. Moreover, the functional loss of *DDOST* in humans, demonstrates a phenotype with characteristics known as congenital disorders of glycosylation (CDG)^28^. A sole case was described in a seven-year-old male child with CDG, whose condition arose from the inheritance of a maternal missense mutation and a paternal point mutation resulting in a premature stop codon and as a result the patient exhibited severe hypoglycosylation^28^. It was shown however that restored wild-type *DDOST* cDNA in the patients’ fibroblasts rescued glycosylation, thus indicating the importance of functional DDOST in a pathological environment^28^.

For the present study, we hypothesised that mice with a global OST48 overexpression would be protected from increases in circulating AGEs and impaired kidney function in the context of diabetes. Specifically, we aimed to identify whether increased OST48 in the presence of hyperglycaemia could drive AGE lowering to protect against AGE-mediated microvascular damage typically seen in DKD, such as glomerular pathology and greater tubulointerstitial fibrosis.

## Results

### Characterisation of diabetes in a site directed global OST48 knock-in mouse model

Mice were generated with a global over-expression of OST48 (*DDOST*+/-)^29^. These mice exhibited variation in food and water consumption, urine output and a significant reduction in kidney weight compared with wild-type mice (**Table 1**). Diabetes was characterised by elevated GHb, fed/fasting blood glucose and increased fasting insulin (**Table 1**). Diabetic mice had significantly lower mean total body weight, increased urine output as well as increased food and water consumption and renal hypertrophy (**Table 1**).

**Table 1.** Biochemical/anthropometric measurements post-diabetes (Week 24 of study/32 weeks of age). Mean and standard deviation for biochemical/anthropometric measurements (n = 5-11). Significance levels were determined by two-way ANOVA, testing the effect of genotype and diabetes. Differences between variables identified by Bonferroni’s post hoc test. Bold P values indicates significant effect of at least < 0.05.

|  | **WT** |  | ***DDOST*+/-** |  | **Two-Way ANOVA** |  |  |
| --- | --- | --- | --- | --- | --- | --- | --- |
|  | Control | Diabetic | Control | Diabetic | Genotype | Diabetes | Interaction |
| GHb<br>(mmol/mol) | 5.58 ±<br>0.36 | 11.75 ±<br>2.41 | 6.54 ±<br>1.21 | 10.39 ±<br>2.46 | 0.36 | **<0.0001** | 0.17 |
| Food<br>consumption<br>(g/24 h) | 0.067 ±<br>0.016 | 0.31 ±<br>0.058 | 0.078 ±<br>0.026 | 0.20 ±<br>0.082 | **0.014** | **<0.0001** | **0.0033** |
| Fed Glucose<br>(mmol/L) | 16.26 ±<br>2.43 | 25.30 ±<br>6.22 | 16.26 ±<br>3.18 | 27.25 ±<br>1.50 | 0.52 | **<0.0001** | 0.52 |
| Fasting<br>Glucose<br>(mmol/L) | 10.79 ±<br>3.02 | 12.98 ±<br>2.51 | 9.70 ±<br>0.98 | 12.35 ±<br>1.91 | 0.36 | **0.015** | 0.81 |
| Fasting Insulin<br>(ng/ml) | 0.98 ±<br>0.61 | 0.33 ±<br>0.07 | 0.97 ±<br>0.32 | 0.71 ±<br>0.15 | 0.22 | **0.0056** | 0.21 |
| Water<br>consumption<br>(ml/24 h) | 0.13 ±<br>0.010 | 1.73 ±<br>0.43 | 0.15 ±<br>0.052 | 0.91 ±<br>0.39 | **0.00030** | **<0.0001** | **0.0002** |
| Urine<br>production<br>(ml/24 h) | 0.025 ±<br>0.012 | 1.37 ±<br>0.43 | 0.019 ±<br>0.012 | 0.77 ±<br>0.33 | **0.0030** | **<0.0001** | **0.0035** |
| Kidney<br>Weight/BW<br>ratio (g) | 11.92 ±<br>2.99 | 19.68 ±<br>0.71 | 10.71 ±<br>1.94 | 16.36 ±<br>1.23 | **0.020** | **<0.0001** | 0.25 |

### Increased glomerular OST48 facilitated AGE urinary excretion in *DDOST+/-* mice

SWATH proteomics in renal cortices enriched for glomerular proteins identified that non-diabetic *DDOST*+/- mice had a significant increase in OST48 protein abundancy (48.39-fold increase) over non-diabetic wild-type mice (**Figure 1A**). There was also a tendency toward increased OST48 protein in glomerular fractions from diabetic *DDOST*+/- mice (*P*=0.07; 14.44-fold increase) compared to diabetic wild-type mice (**Figure 1A**). Surprisingly, there was no significant increase in OST48 seen in tubular enriched protein fractions (**Figure 1B)** from *DDOST+/-* mice (**Figure 1C**). Glomerular OST48 appeared to be predominately expressed by podocytes as indicated by the co-localisation of nephrin (**Figure 1D**) as well as within proximal tubule cells expressing SGLT2 (**Figure 1E**). Subsequently, there was a significant increase in renal AGE excretion by all diabetic mice as well as the non-diabetic *DDOST+/-* mice (**Figure 1F**). Specifically, urinary AGE excretion determined by total *N*-(carboxymethyl)lysine (CML) was increased by ∼120% in non-diabetic DDOST+/- mice compared to wild-type mice (**Figure 1F**; *P*=0.033). However, *DDOST+/-* mice showed no increases in AGE accumulation in either kidney tissue (**Figure 1G**) or within the circulation (**Figure 1H**).

**Figure 1.**
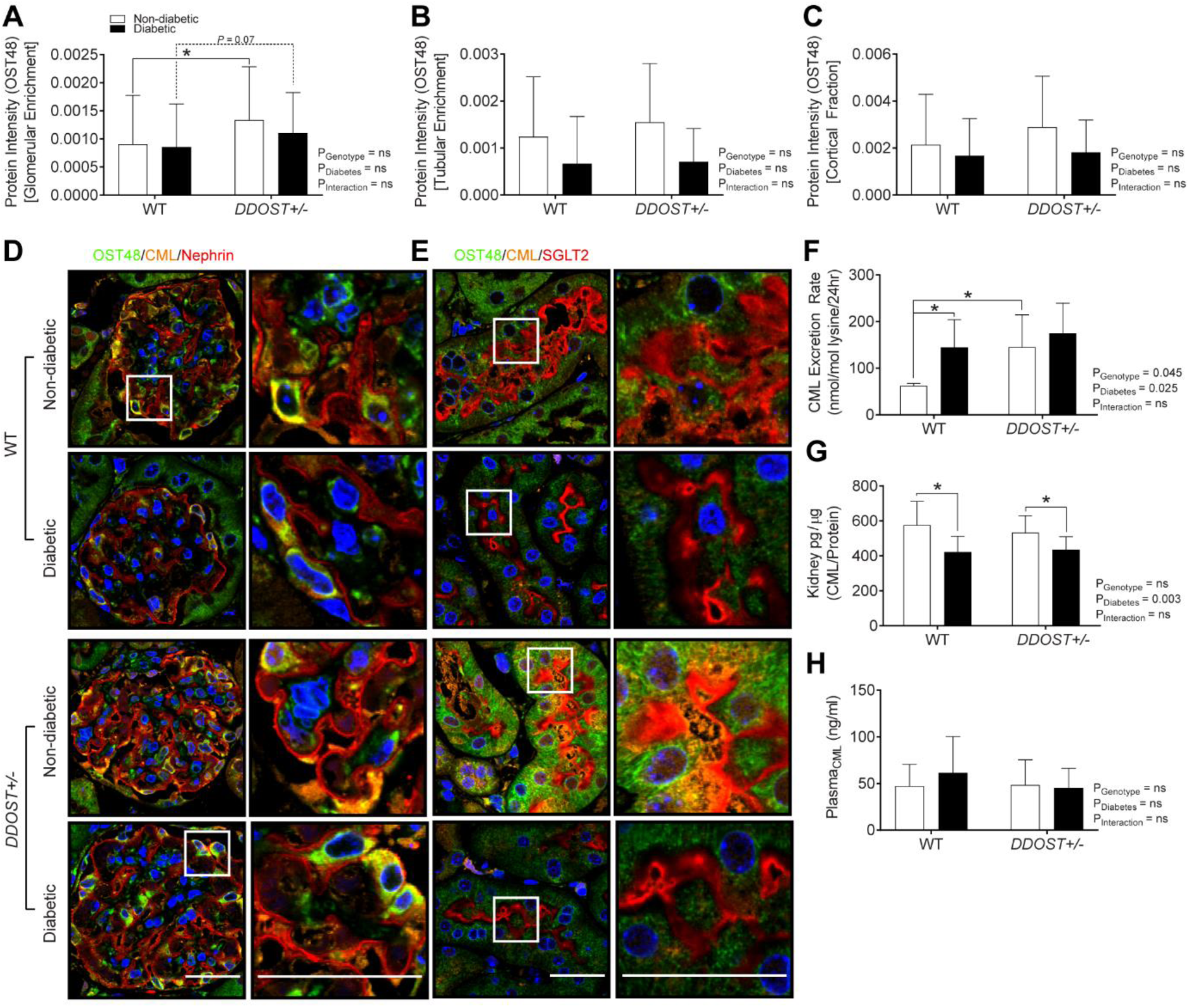
*DDOST+/-* mice have increased glomerular expression of OST48, increased CML excretion, however no changes in CML deposition in the kidney or plasma CML levels. OST48 expression and localization in the kidney. SWATH proteomics (**A-C**) quantification of the protein intensities of OST48 in processed kidney tissue enriched for either (**A**) glomeruli or (**B**) tubules and in the kidney cortex (**C**). Confocal photomicrographs (**D-E**) of OST48 (green), CML (orange) and either a podocyte foot process marker, nephrin (red) or a proximal tubule marker, SGLT2 (red) on kidney sections imaged at either a (**D**) glomerulus or (**E**) proximal tubule. CML ELISA (**F-H**) measuring the total content of CML in (**F**) 24-hour urine collections, (**G**) whole kidney and (**H**) plasma. Results are expressed as mean ± SD with either two-way ANOVA or unpaired t-test analysis (n = 4-9) *P < 0.05. Scale bars from representative images for confocal microscopy are representative of 5µm. For proteomics, MSstatsV3.10.0 determined significant (P < 0.05) log fold changes in the protein intensities between the selected experimental group the wild-type non-diabetic group (n = 4-5).

### Diabetic *DDOST*+/- mice had no improvements in kidney function despite increased urinary AGE excretion

All diabetic mice had impaired kidney function as ascertained by a decrease in creatinine clearance **(Figure 2A)**, which was not affected by OST48 overexpression. Diabetic mice also had albuminuria when assessed by either albumin:creatinine ratio (**Figure 2B**) or 24 hour urinary albumin excretion (**Figure 2C**). Therefore, increased OST48 expression did not prevent the decline in kidney function which is characteristic of diabetic kidney disease.

**Figure 2.**
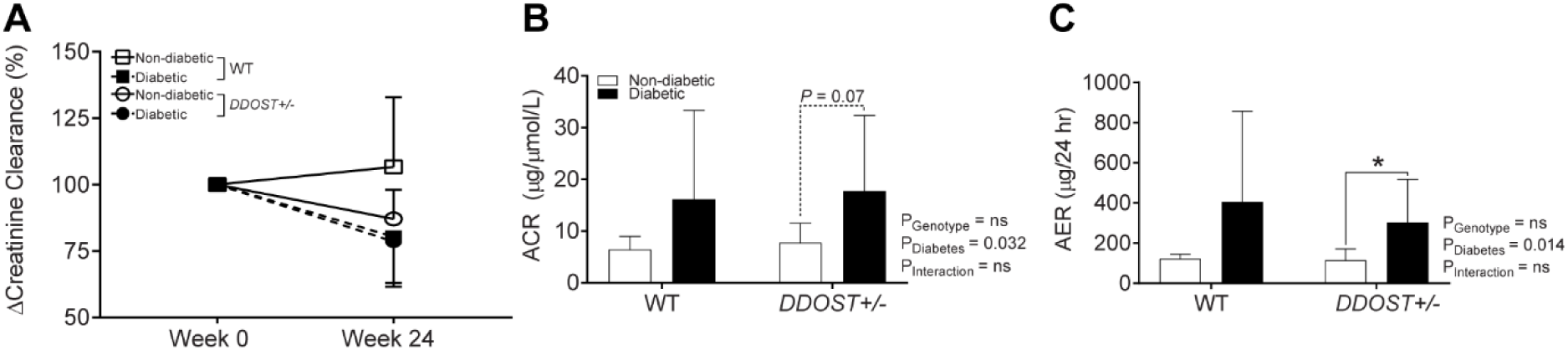
Diabetic mice with global increases in OST48 are not protected against the development of diabetic kidney disease. Serum and 24-hour urine was collected from mice in metabolic cages at week 0 and week 24 of the study. Creatinine was measured spectrophotometrically and the (**A**) change in creatinine clearance over the study period was determined from matched creatinine clearance values from week 0 of the study (6-8 weeks of age) to week 24 of the study. (**B**) Urinary albumin:creatinine ratio measured at week 24 of the study. (**C**) Albumin was measured spectrophotometrically at 620nm in a biochemical analyzer and the albumin excretion rate (AER) was determined based on the 24-hour urine flow rate. Results are expressed as mean ± SD with either two-way ANOVA or paired t-test analysis (n = 3-8) *P < 0.05.

### Globally increasing OST48 expression did not protect against diabetes-induced kidney structural damage

Diabetes resulted in glomerulosclerosis in both genotypes to the same degree by the study end (**Figure 3A**). This was supported by SWATH proteomics data from kidney cortices enriched for glomerular proteins which demonstrated significant increases in the abundance of collagen proteins in diabetic mice (**Figure 3B**). Increasing the expression of OST48 also did not prevent tubulointerstitial fibrosis (TIF) in the kidneys of diabetic mice, as seen with Masson’s trichrome (**Figure 3C**) and Sirius red (**Figure 3D**) staining. Indeed, there was exacerbation of TIF in diabetic *DDOST+/-* by comparison to diabetic wild-type littermates (**Figure 3D**; 32.0% increase, *P=*0.043).

**Figure 3.**
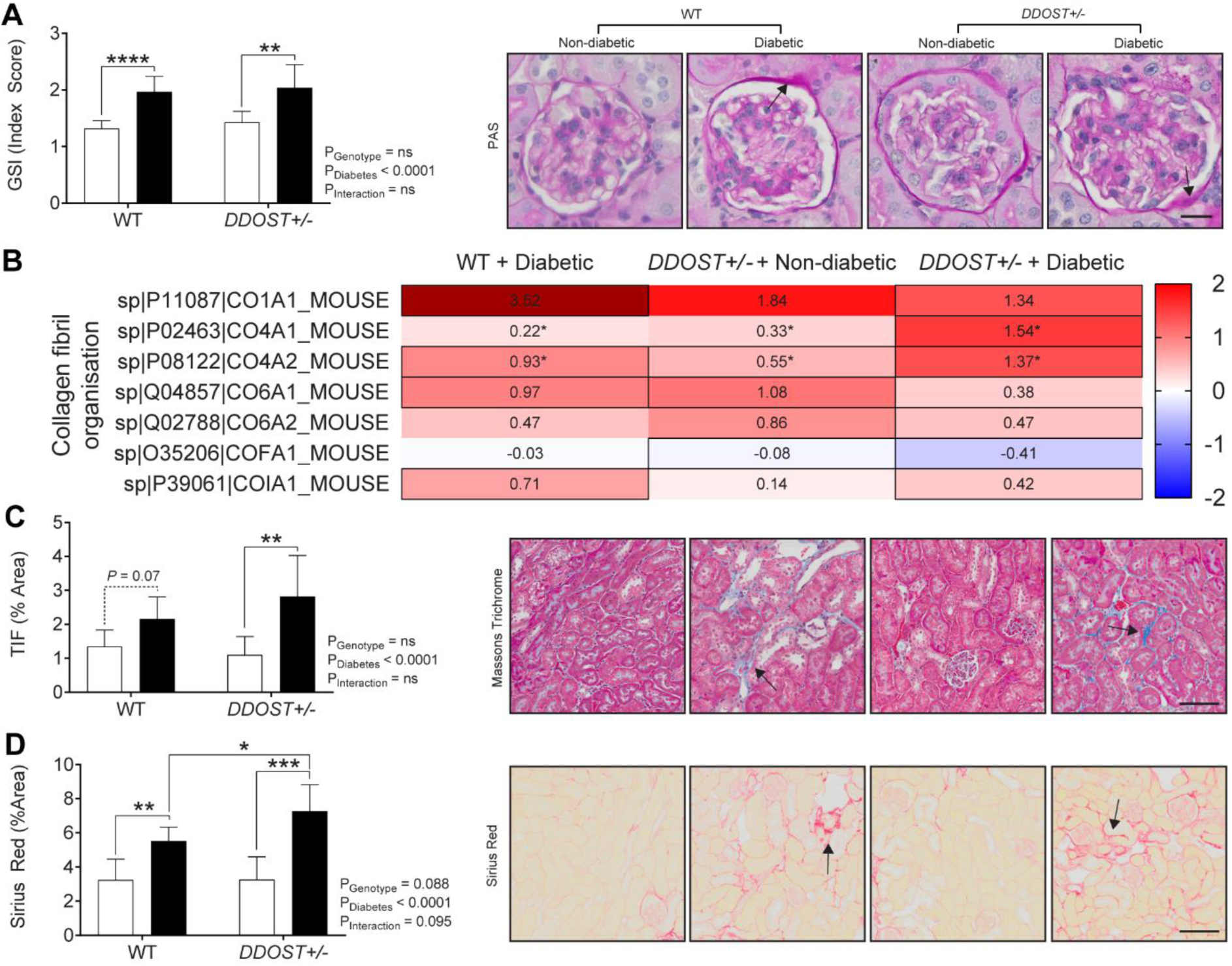
DDOST+/- mice were not protected from renal structural damage and exhibited similar patterns of collagen protein content exhibited in mice with diabetes. (**A**) Assessment of renal glomerular damage in the kidney with Periodic-acid Schiff staining (PAS), which was then quantified based on a positive threshold protocol. *DDOST+/-* mice and mice with a diabetic phenotype exhibited moderate glomerulosclerosis, indicated by an increase in mesangial matrix expansion (black arrow). (**B**) Heat map representation of SWATH-MS proteomics of glomeruli enriched proteins for enzymatic pathways involved in collagen fibril organization. Significant proteins are represented as bolded cells, where red indicates an increase and blue indicates a decrease in protein concentrations. (**C**) Presence of collagen in Masson’s trichrome (blue staining) and (**D**) Sirius red (red staining) in the interstitium of the tubules (black arrow) is an indicator of progressive kidney damage. The severity of these changes was more pronounced in mice with a diabetic phenotype. Scale bars from representative images of (**A**) glomeruli stained with PAS or (**C-D**) tubule sections stained with either Masson’s trichrome or Sirius red were 20µm and 100µm, respectively. Results are expressed as mean ± SD with either two-way ANOVA or unpaired t-test analysis (n = 5-9) *P < 0.05, **P < 0.01, ***P < 0.001, ****P < 0.0001. For proteomics, MSstatsV3.10.0 determined significant (P < 0.05) log fold changes in the protein intensities between the selected experimental group the wild-type non-diabetic group (n = 3-5).

### Diabetic mice with globally increased OST48 expression have greater insulin secretory capacity and increased P13K-AKT activity in glomeruli, which is dissociated from deterioration in kidney function

In the absence of diabetes, globally increasing the expression of OST48 had no significant effect on insulin sensitivity or glucose tolerance (**Figure 4A-G**). However, when diabetes was induced, mice with a global over-expression of OST48 had significant reduction in fractional glucose excretion (**Figure 4A**), increases in fasting plasma insulin (**Figure 4B**) and lower blood glucose concentrations during an ipGTT, both acutely at 15 minutes (11.9% decrease, *P* = 0.097) and 30 minutes (14.2% decrease, *P =* 0.017) post-bolus of D-glucose (**Figure 4C**). Consistent with these changes, overall insulin secretion (**Figure 4D;** average 77.4% increase, *P* < 0.05 – 0.001), first phase insulin secretion (**Figure 4E**, 99.5% increase, *P* = 0.0005) and the AUC insulin (**Figure 4F**) during the ipGTT were all greater in diabetic mice over expressing OST48, when compared to littermate wild type diabetic mice. Insulin effectiveness (AUIC:AUGC) was also greater in diabetic *DDOST+/-* mice compared to diabetic wild-type mice (**Figure 4E-G**; 99.5% increase, *P*=0.0005). During fed conditions an insulin tolerance test (ipITT), all diabetic *DDOST*+/- mice had decreased responsiveness to insulin with persistently elevated plasma glucose concentrations, when compared to non-diabetic mice (**Figure 4H**) but did not differ between genotypes. This was particularly evident during the first 60-minutes of the ipITT, where there was a rapid decline in plasma glucose concentrations in non-diabetic mice (**Figure 4H-I**; 490% decline compared to diabetic mice), with rapid recovery of blood glucose concentrations by 120 mins. This increase in insulin secretion and effectiveness was also confirmed by increased PI3K-AKT activation in the glomerular enriched fractions from mice over-expressing OST48 **Figure 4J**).

**Figure 4.**
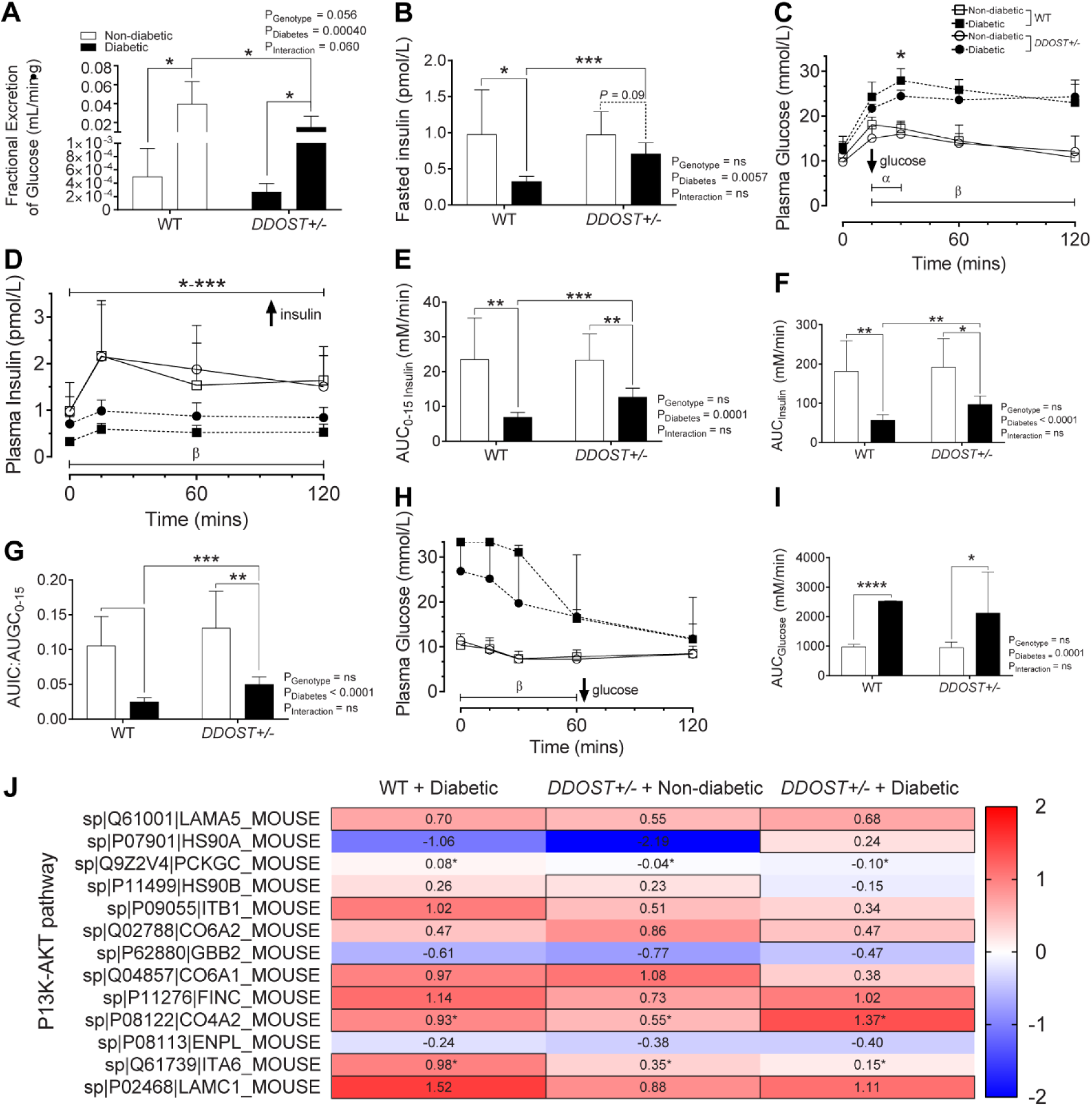
Diabetic *DDOST+/-* mice exhibit increased first phase insulin secretion and activation of the P13K-AKT pathway akin to a diabetic phenotype. (**A**) 24-hour fractional excretion of glucose. (**B**) Fasting plasma insulin concentrations. (**C**) Plasma glucose and (**D**) insulin curve over 120-minutes following an intraperitoneal glucose injection and (**E**) first phase area-under-the-curve (AUC) analysis and (**F**) 120-minute AUC analysis. (**G**) Ratio of AUIC:AUGC of early-phase insulin response. (**H**) Plasma glucose over 120-minutes following an intraperitoneal insulin injection and (**I**) 120-minute AUC analysis. (**J**) Heat map representation of SWATH-MS proteomics data for enzymatic pathways in glomerular proteins involved the P13K-AKT signalling pathway. Significant proteins are represented as the Log2 fold change where red indicates a decreased and blue indicates an increase in protein concentrations. Results are expressed as mean ± SD with either two-way ANOVA or unpaired t-test analysis (n = 5-9) \**P <* 0.05, \*\**P*<0.01, \*\*\**P*<0.001. For IPGTT and IPITT curves results are represented as \**P*<0.05, \*\**P<*0.01, \*\*\**P*<0.001 for *DDOST+/-* diabetic compared to wild-type diabetic, α < 0.05 effect due to genotype, β < 0.05 effect due to diabetes. For proteomics, MSstatsV3.5.1 determined significant (P < 0.05) log fold changes in the protein intensities between the selected experimental group the wild-type non-diabetic group (n = 3-5).

## Discussion

There has been some speculation that increasing AGE clearance via increases in OST48 could alleviate the development and progression of diabetic kidney disease^19,20,22,30,31^. We have shown for the first time that globally increasing OST48 through the over-expression of *DDOST* in a mouse model of diabetes, did not protect against kidney disease. Indeed, there was an exacerbation of tubulointerstitial fibrosis seen in diabetic mice with a global over-expression of OST48. Quite surprisingly, however, this was despite significant improvements in insulin secretory capacity and action in these diabetic *DDOST+/-* mice. Therefore, this study **dissociated improvements in insulin secretion from slowing the onset and progression of DKD**, in mice over-expressing OST48.

Previous studies have suggested that increasing AGE clearance from the body through the OST48 pathway prevents oxidative stress^19^ and inflammatory responses^32^, as well as improving healing and hyperglycaemia^26^, concluding that increased OST48 activity could have potential benefits to alleviate vascular complications in diabetes^19,31^ including DKD^19,32^. Moreover, previous studies have alluded at the potential benefits of AGE restriction and reducing serum AGE levels in diabetic patients^24,33-35^, and suggested that AGE excretion facilitated through increased OST48 could reduce the AGE burden^16,24,26,36,37^. We have recently published data evidence of a global overexpression of OST48 in combination with a secondary impact of a high AGE diet causes liver abnormalities which likely contribute to changes in insulin handling and a metabolic shift of fuel utilisation^29^. We were surprised, that increasing AGE clearance by the kidneys, via globally increasing OST48 expression had no effect on kidney function and structural parameters in this mouse model of diabetes and worsened renal pathology.

In our mouse model the lack of effect of OST48 to alleviate DKD was despite modest improvements to insulin secretory capacity, although this was not sufficient to improve the long-term markers of glycaemic control (GHb), nor fasting and fed plasma glucose concentrations. However, our data is in agreement with the DCCT/EDIC clinical trials, which indicate that without effective reduction in blood glucose early in disease development, there was no reduction in the risk in development of diabetic vascular complications including DKD^38,39^. There have been previous studies where AGEs have been shown to impair insulin secretion^40,41^ although it is unclear as to the role that OST48 plays in beta cells. However, we have identified that OST48 is expressed in beta cells^42^. In addition, further investigation would be required to elucidate why these apparent improvements in insulin secretion seen in diabetic OST48 mice did not translate to improved long-term clinical markers of glycaemic control. However, this may just indicate that modest improvements in insulin secretion later in disease are not sufficient to alter the progression of diabetic kidney disease.

There was also a decline in renal AGE content seen with diabetes, which was consistent with the increases in AGE excretion into the urine, but this was not affected by OST48. In agreement with our findings, another study showed that increasing OST48 expression in mice also facilitated urinary AGE clearance, without affecting renal AGE concentrations^26^. This would suggest that rather than specific AGE uptake into the kidney the flux of AGEs through the kidney into the urine, may trip signalling cascades for other AGE receptors such as RAGE^43^. This could be feasibly responsible for the observed glomerulosclerosis and tubulointerstitial fibrosis^44^ with diabetes and with OST48 overexpression where chronic binding of circulating AGEs to RAGE results in diabetic kidney and cardiovascular disease^35,44,45^. Increased AGE flux from the circulation to the urine, is also postulated in other studies to have detrimental effects on proximal tubules contributing to tubulointerstitial fibrosis^46^, which is consistent with our model. However, we do not suspect defects in protein N-glycosylation contributed to the kidney pathology seen with OST48 overexpression ^47^. This is because, we have previously shown that ubiquitous global overexpression of OST48 does not impair nor alter protein N-glycosylation^29^.

Despite the ineffectiveness of increased OST48 to protect against the development of diabetic kidney disease, we did however observe benefits on insulin secretion, fasting insulin and insulin sensitivity by globally elevating OST48 in diabetic mice. Our data suggests that downstream insulin receptor activity via P13K and diverging into increased lipogenesis, glycolysis and glycogenesis (starch and sucrose metabolism) occurs in the kidney when OST48 is increased. Our data is in concordance with previous human studies which suggest that AGE restriction whether through dietary^24^ or therapeutic^33^ methods improve insulin sensitivity and decrease in serum OST48 concentrations^24^. We have also consistently shown that AGE lowering therapies which decrease the burden of AGEs improve both insulin secretion^40^ and improve insulin sensitivity^48,49^.

In summary, these studies revealed that increasing OST48 expression globally does not prevent the development of DKD. Specifically, in diabetic mice, increasing OST48 expression did not prevent the decline in kidney function, glomerulosclerosis and exacerbated tubulointerstitial fibrosis. Therefore, we would suggest that increasing the flux of AGEs into the urine is not a strategy worth pursuing in DKD. Similarly, it appears that increasing AGE signalling in the kidney via receptors such as RAGE without increasing kidney AGE content, is responsible for this lack of protective effect. Therefore, this study suggests that globally increasing the expression of OST48 thereby increasing urinary AGE excretion does not improve kidney function in the context of diabetes. However, further studies are required to identify whether kidney cell-specific modulation of OST48 may be superior to a global approach.

## Materials and Methods

### Animal husbandry

Male C57BL/6J wild-type mice and littermate heterozygotes with a ubiquitous genetic insertion of the human gene encoding AGE-R1/OST48 (*DDOST*+/-) at the ROSA26 locus (*ROSA26*^*tm1* (*DDOST*)*Jfo*^; termed as *DDOST+/-*) were generated (Ozgene, Australia). Between 6-8 weeks of age (Week 0), diabetes was induced in male wildtype (n = 9) and *DDOST+/-* (n = 9) mice by multiple low dose intraperitoneal injections of streptozotocin (STZ)^50^, at a dosage of 55mg/kg/day (dissolved in sodium citrate buffer, pH 4.5) for 5 consecutive days. Control or non-diabetic male wild-type (n = 11) or *DDOST+/-* (n = 12) mice received equivalent injections of sodium citrate buffer alone. After 10 days of recovery, blood glucose concentrations were determined and then repeated weekly to ensure mice included were diabetic (blood glucose concentrations > 15mmol/L). Mice were housed in specific pathogen free housing conditions and allowed access to food and water ad libitum and were maintained on a 12 hour light:dark cycle at 22°C. All mice received a diet of standard mouse chow low in AGE content (AIN-93G^40^; Specialty Feeds, Australia) *ad libitum*. At 0 and 12 weeks of the study, mice were housed in metabolic cages (Iffa Credo, l’Arbresle, France) for 24 hours to determine food and water consumption. Urine output was also measured during caging and urinary glucose determined by a glucometer (SensoCard Plus, POCD, Australia).

### Intraperitoneal glucose and insulin tolerance testing (IPGTT/IPITT)

Intraperitoneal glucose (ipGTT) and insulin (ipITT) tolerance tests were performed as outlined previously^51^. Briefly, for ipGTT experiments, mice were fasted for 6 hours and a 1g/kg bolus of glucose was injected intraperitoneally (ip) and 50µl of blood was sampled at 0, 15, 30, 60, and 120 mins for plasma glucose and insulin analysis. During ipITT experiments, mice were injected with 1.0U of fast acting insulin/kg (Humulin) diluted in 0.9% saline and 50µl of blood was sampled at 0, 30, 60, and 120 mins for plasma glucose analysis.

### Liquid Chromatography-Mass Spectrometry (LC-MS/MS)

As previously described^52^, proteins were extracted from whole liver tissue samples using guanidine denaturing buffer (6 M guanidinium, 10 mM DTT and 50 mM Tris-HCl). Reduced cysteines were alkylated with acrylamide, and quenched with excess DTT. Proteins were precipitated in 4 volumes of 1:1 methanol:acetone and digested with trypsin. Peptides were desalted and analysed by Information Dependent Acquisition LC-MS/MS as described^53^ using a Prominence nanoLC system (Shimadzu, NSW, Australia) and Triple TOF 5600 mass spectrometer with a Nanospray III interface (SCIEX). SWATH-MS analysis was performed^54^ and analysed with MSstats as previously described. Differentially abundant proteins were analysed using DAVID^55^.

### Serum/urine creatinine and clearance

Serum and urinary creatinine were measured spectrophotometrically at 550nm (Cobas Mira, Roche Diagnostics, Australia), and the ratio of creatinine clearance was determined as specific measure of renal function.

### Albumin excretion rate

Urine albumin was measured spectrophotometrically at 620nm (FLUOstar Optima, BMG Labtech, Australia), and 24-hour albumin excretion rate was calculated by normalizing the levels of albumin to the flow rate of urine.

### Histology and imaging

Paraffin-embedded sections were stained with either a Periodic acid Schiff (PAS) staining kit (Sigma-Aldrich, United States), a Trichrome (Masson) staining kit (Sigma-Aldrich, St. Louis, Missouri, USA) or Sirius Red (Sigma-Aldrich, United States). All sections were visualized on an Olympus Slide scanner VS120 (Olympus, Japan) and viewed in the supplied program (OlyVIA Build 10555, Olympus, Japan). Slides were quantified based on threshold analysis in Fiji^56^. Briefly, for immunofluorescence staining, paraffin-embedded sections were stained with a combination of either anti-CML (1:200 dilution; ab27684; Abcam, United Kingdom), OST48 (H-1; 1:100 dilution; SC-74408; Santa Cruz biotechnologies, United States), nephrin (1:100 dilution; ab27684; Abcam, United Kingdom) and SGLT2 (M-17; 1:100 dilution; sc-47403; Santa Cruz biotechnologies, United States). Confocal images were visualized on an Olympus FV1200 confocal microscope (Olympus, Japan) and viewed in the supplied program (FV10, Olympus, Japan).

### Glomerulosclerotic index (GSI)

GSI as a measure of glomerular fibrosis was evaluated in a blinded manner by a semi-quantitative method^57^. Severity of glomerular damage was assessed on the following parameters; mesangial matrix expansion and/or hyalinosis of focal adhesions, true glomerular tuft occlusion, sclerosis and capillary dilation. Specifically, grade 0 indicates a normal glomerulus; 1, <25% glomerular injury; grade 2, 26-50%; grade 3, 51-75%; and grade 4, >75%.

### Statistical Analyses

Results are expressed as mean ± SD (standard deviation), and assessed in GraphPad Prism V7.01 for Windows (GraphPad Software, United States). Normally distributed parameters (tested with D’Agostino & Pearson omnibus normality test) were tested for statistical significance by 2-way ANOVA followed by post hoc testing for multiple comparisons using the Bonferroni method unless otherwise specified. For comparison between groups as required, a two-tailed unpaired Student’s t-test was used where specified. For SWATH-MS, MSstatsV3.10.0^58^ was used to detect differentially abundant proteins estimating the log-fold changes between compared conditions of the chosen experimental group and with the wild-type non-diabetic mice. For all calculations, a P < 0.05 was considered as statistically significant.

### Experimental animal ethics statement

All animal studies and experiments were performed in accordance with guidelines provided and approved by the AMREP (Alfred Medical Research and Education Precinct) Animal Ethics Committee and the National Health and Medical Research Council of Australia (E/0846/2009B).

### Data availability statement

The datasets generated and analysed during the current study are available from the corresponding author on reasonable request.

## Acknowledgements

The authors would like to acknowledge Maryann Arnstein and Anna Gasser at Baker IDI Heart and Diabetes Institute, Australia, and David Briskey at the School of Human Movement and Nutrition Sciences, University Queensland, Australia, for technical assistance.

## Author Contributions

The following authors’ contributions are listed below. AZ, JMF and BS wrote the main manuscript text, AZ prepared figures 1-5 and supplementary figures, in addition to the accompanied figure legends and supplementary figure legends. FYTY, DM, CL, SSL, KCS, BEH, SAP, VTB and MTC assisted with experiments and/or revised the manuscript.

